# CTFacTomo: Reconstructing 3D Spatial Structures of RNA Tomography Transcriptomes by Collapsed Tensor Factorization

**DOI:** 10.1101/2024.11.11.623086

**Authors:** Tianci Song, Quoc Nguyen, Charles Broadbent, Rui Kuang

**Affiliations:** Department of Computer Science and Engineering, University of Minnesota Twin Cities, Minneapolis, 55414, MN, USA

**Keywords:** Spatial Genomics, RNA Tomography Sequencing, Collapsed Tensor Factorization

## Abstract

Cells are organized to form three-dimensional structures of complex tissues. To map the complete 3D organization of a tissue, technologies based on tissue microdissections provide deep bulk RNA sequencing of orthographically arranged cryosections of a tissue, such that the full 3D spatial structure could be inferred from deeply sequenced transcriptomes in three views projected similarly as 3D tomography. Here, we introduce CTFacTomo to learn a Collapsed Tensor Factorization for RNA tomography data from cryosections to reconstruct 3D spatially resolved gene expressions. CTFacTomo combines tensor factorization with collapsing tensor entries to match the bulk gene expressions in each cryosection, enriched by a regularization of a product graph of protein-protein interaction network and spatial graphs. In the experiments, CTFacTomo is first validated on three datasets projected from fully profiled 3D spatial gene expressions to demonstrate that CTFacTomo significantly outperforms the benchmark methods for predicting the ground-truth gene expressions based on the projected 1D spatial gene expressions of three orthographic views. CTFacTomo is then applied to two RNA tomography datasets from zebrafish embryo and mouse olfactory mucosa, respectively. In both datasets, CTFacTomo detects 3D spatial expressions of several marker genes that are consistent with the developmental or functional regions in comparison to accompanying ISH staining images. In addition, a qualitative comparison between the reconstructed zebrafish embryo gene expressions with a matched external 3D Stereo-seq dataset also suggests that CTFacTomo reconstructs more spatially coherent patterns in the whole transcriptome with state-of-the-art performance.

## 1 Background

Cells in multicellular organisms are organized to form complex 3D tissue structures and create distinct microenvironments to drive essential events such as cell movement, cell communication and interactions, and spatially varying gene expressions. For example, morphogen gradients diffuse through 3D space to regulate cell differentiation and morphogenesis, orchestrating the development of functional organs during embryogenesis. To reveal the spatial organization of the cells and their functions in the tissues, spatial transcriptomic technologies such as in-situ hybridization (ISH) (Raj et al. 2008; Chen et al. 2015; Lubeck et al. 2014; Shah et al. 2016; Eng et al. 2019; Lee et al. 2014; Wang et al. 2018) or in-situ capturing (ISC) (Ståhl et al. 2016; Rodriques et al. 2019; Vickovic et al. 2019; Chen et al. 2022) have been widely used to profile gene expression with retained spatial localization information in the tissue (Moses and Pachter 2022). Among these spatial profiling technologies, technologies based on microdissection adopt a different strategy, which isolates slices or regions of tissues for separate RNA sequencing such that the location of the cells is informed by the organization of the arranged slices or regions (Li and Peng 2022). Transcriptome tomography sequencing (Tomo-seq) (Junker et al. 2014) is a notable method in this category inspired by image tomography. In Tomo-seq, replicates of a 3D tissue are dissected into successive slices along three orthogonal views (axes), and then the RNAs in the cells in each slice are collected for deep bulk RNA sequencing. Finally, the 3D structures of the gene expressions could be reconstructed by the gene expressions measured from the slices stacked in each of the three axes using a computational method.

Tomo-seq provides more reliable transcriptome-wide gene expression measurements through highly sensitive bulk RNA sequencing, effectively avoiding key limitations, namely that ISC-based methods generally suffer from low RNA capture rates, while ISH-based methods are limited by lower detection sensitivity in transcriptome-wide profiling. More importantly, since both ISH and ISC inherently profile RNAs in a 2D space, a challenging step is required to align stacked slices by a manual-intensive registration of annotated regions, tissue histological images, or spatial coordinates for 3D construction. In contrast, Tomo-seq performs RNA sequencing of slices dissected in three views from replicates for a real 3D measuring of the gene expressions for full 3D reconstruction. In addition, the cost for library preparation and sequencing of cryosection samples are significantly lower than using fluorescent-imaging or spatial-array based profiling of 3D tissues in typical experiments. Despite these advantages, one significant limitation of Tomo-seq and other microdissection-based methods is that the reconstruction of the 3D spatial gene expression is a computational challenge and setback to a great extent. The current solution for 3D reconstruction with Tomo-seq is the iterative proportional fitting (IPF) algorithm (Fienberg 1970), which is a relatively simplistic heuristic optimization method for minimizing fitting error for each individual gene independently. Accordingly, IPF is prone to noise in RNA sequencing data and might fail to capture complex spatial patterns (Schede et al. 2021).

In this paper, we introduce Collapsed Tensor Factorization for Tomography (CTFacTomo) to learn a tensor factorization representation of spatially resolved 3D gene expressions from RNA tomography data of cryosections. CTFacTomo reconstructs 3D spatial transcriptomics based on the 1D bulk RNA sequencing data obtained for the consecutive slices in each orthogonal spatial axis of tomography along the tissue as outlined in Fig. 1. The 1D gene expression data represent the expression of each gene in each consecutive slice in one orthogonal axis (Fig. 1a). Then, the 1D data in three views is combined to reconstruct the 3D spatial transcriptomes in a 4-way tensor with x-y-z spatial modes and gene mode as shown in Fig. 1b. CTFacTomo combines tensor factorization with a loss function supervised by collapsing the tensor entries to match the gene expressions in each cryosection and the 3D tissue mask. CTFacTomo outputs a Canonical Polyadic decomposition of the reconstructed tensor aligned with the spatial axes. CTFacTomo also incorporates spatial relations among the slices along different axes and gene functional relations among genes with regularization by a product of a protein-protein interaction (PPI) network and spatial graphs to utilize important prior information (Fig. 1b). 3D spatial expressions can be reconstructed from the decomposition model for identifications of 3D spatial patterns of marker genes and other spatial characteristics of the transcriptome. (Fig. 1c).

**Fig. 1.**
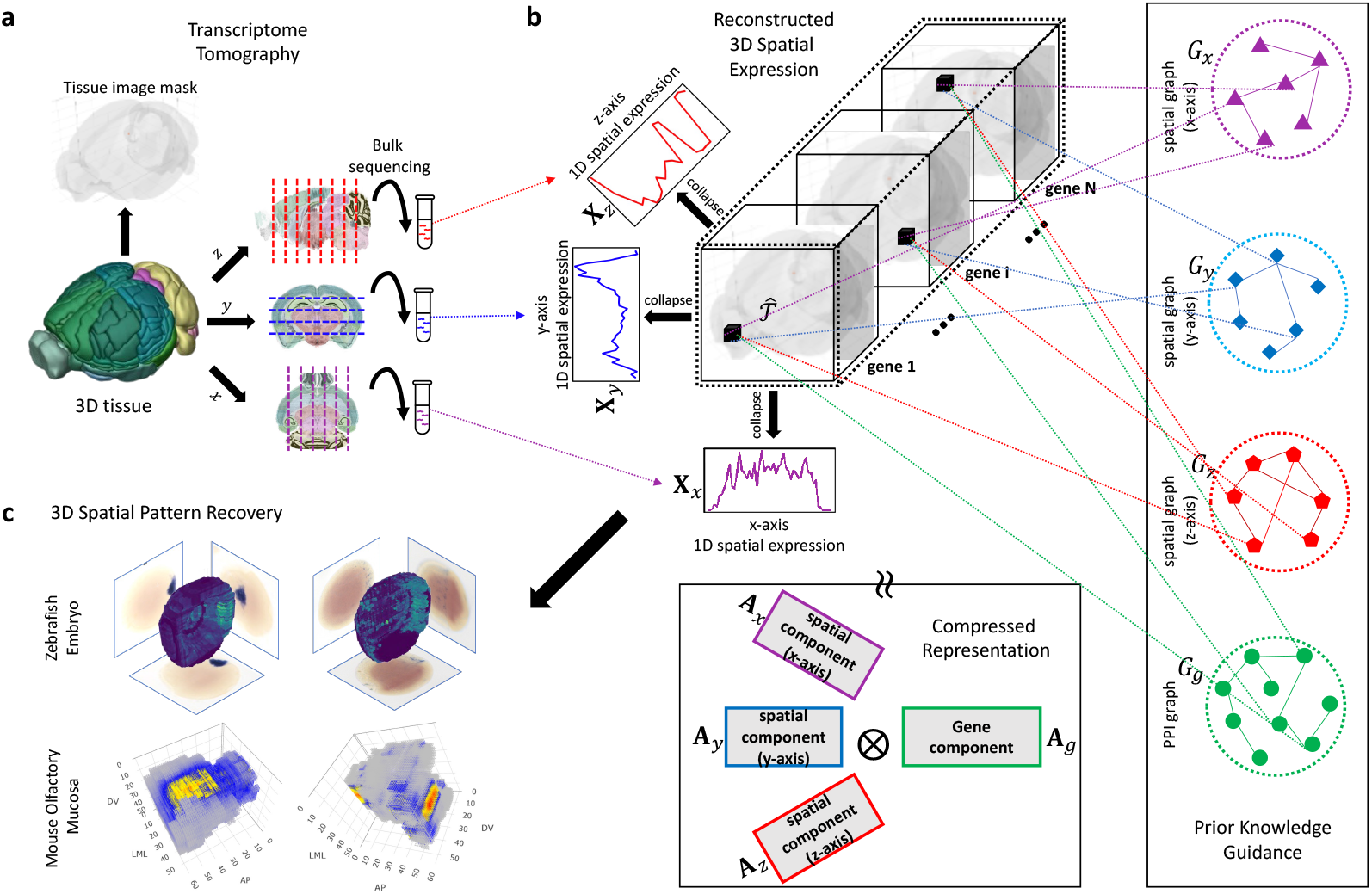
Overview of CTFacTomo. **a** Transcriptome tomography generates high-coverage 1D spatial transcriptomics along three orthogonal spatial axes. A 3D tissue and its replicates are sectioned into successive slices along each axis, and bulk transcriptomics techniques are applied to quantify gene expression along the axis. Typically, a 3D mask of the profiled tissue can also be constructed from imaging. **b** Given 1D spatial gene expression of each gene along three orthogonal spatial axes (**X**_*x*_, **X**_*y*_, **X**_*z*_), CTFacTomo reconstructs 3D spatial gene expression as a 4-way tensor 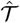. The reconstruction is estimated via CP decomposition with four factor matrices (**A**_*x*_, **A**_*y*_, **A**_*z*_, **A**_*g*_), minimizing the reconstruction error with respect to the observed projections and the 3D tissue mask. CTFacTomo also incorporates prior knowledge from spatial graphs (*G*_*x*_, *G*_*y*_, *G*_*z*_) and a protein-protein interaction (PPI) network *G*_*g*_ to guide the 3D expression reconstruction. **c** The reconstructed 3D spatial expression enables identifications of 3D spatial patterns of marker genes and other spatial characteristics of the transcriptome. Examples are shown for two large-scale 3D Tomo-seq datasets, one of zebrafish embryo and the other of mouse olfactory mucosa.

In the experiments, CTFacTomo is first validated on three datasets projected from fully profiled 3D spatial gene expressions to demonstrate that CTFacTomo can accurately predict the ground-truth gene expressions based on the projected 1D spatial gene expressions of three orthographic views. Then, CTFacTomo was applied to reconstruct 3D spatially resolved gene expressions in two large-scale 3D RNA tomography datasets, one from a zebrafish embryo and the other from a mouse olfactory mucosa, as shown in Fig. 1c. The reconstructed spatial expressions are evaluated by comparisons with the accompanying ISH staining images of marker genes. Finally, an external Stereo-seq dataset of zebrafish embryo is also matched and used for the evaluation of the reconstructed gene expressions from the RNA tomography data.

## 2 Results

Three experiments were conducted for the evaluation of CTFacTomo. The first experiment focused on three pseudo RNA tomography datasets constructed from projections of 3D spatial transcriptomics data, including two low-resolution ST1K datasets: a mouse brain tissue with 40 slices Ortiz et al. (2020) and a human heart tissue with 9 slices Asp et al. (2019), and a high-resolution Stereo-seq dataset from a drosophila embryo with 21 slices (Wang et al. 2022). The second experiment was conducted on two real RNA tomography datasets, one from a zebrafish embryo ((50, 49, 56) slices) Junker et al. (2014) and the other from a mouse olfactory mucosa ((56, 54, 60) slices) Segura et al. (2022). Finally, for both qualitative and quantitative evaluation, the reconstructed data of the zebrafish embryo was also compared with a 3D Stereo-seq dataset with the matched stage and tissue region Liu et al. (2022). All these datasets are summarized in Table 1. The detailed processing of the datasets is described in the **Methods** section.

**Table 1.** Summary and specifications of the datasets.

| Experiment | Tissue | Platform | Original Size<br>(genes,x,y,z) | Reconstructed Size<br>(genes,x,y,z) |
| --- | --- | --- | --- | --- |
| **3D Projection**<br>**Validation** | Mouse adult brain | ST1K | $10,239 \times 35 \times 33 \times 30$ | $10,239 \times 30 \times 30 \times 30$ |
| | Developing human heart | ST1K | $14,726 \times 35 \times 33 \times 9$ | $14,726 \times 20 \times 20 \times 9$ |
| | Drosophila embryo | Stereo-seq | $14,270 \times 124 \times 105 \times 21$ | $14,270 \times 63 \times 53 \times 21$ |
| **Tomography**<br>**Reconstruction** | Mouse olfactory mucosa | Tomo-seq | $6,153 \times (50 + 49 + 56)$ | $6,153 \times 50 \times 49 \times 56$ |
| | Zebrafish Embryo | Tomo-seq | $9,254 \times (56 + 54 + 60)$ | $9,254 \times 56 \times 54 \times 60$ |
| **External**<br>**Evaluation** | Zebrafish Embryo | Stereo-seq | $5,934 \times 6064(xy) \times 12$ | - |

### 2.1 CTFacTomo accurately reconstructs 3D spatial gene expressions from projected 3D spatial transcriptomics data

The task in this experiment is to reconstruct the ground-truth 3D expressions based on the tomographically aggregated gene expressions from the three curated 3D spatial transcriptomics datasets listed in Table 1. The spatial gene expressions in the 3D datasets were regarded as ground truth, and the 3D expressions were projected into three 1D spatial gene expression matrices that resemble RNA tomography data along the three orthogonal spatial x-y-z axes, each matching a dimension of the 3D spatial expression tensor.

In this experiment on the projected 3D ST data, hyperparameters {*α, β*, rank} were set as {1e2, 1e−3, 120} for the mouse brain dataset, {1e4, 1e−2, 96} for human heart dataset and {1e2, 1e−3, 120} for drosohpila embryo data for reconstruction using the parameter tuning strategy described in the **Methods** section. The complete results for hyperparameter tuning are shown in Fig. 2a-c. The reconstruction performance on the projected ST data was evaluated by mean squared error (MSE) and mean absolute error (MAE) between the reconstructed expression tensor and the ground-truth ST expressions. We also measured both spot-wise and gene-wise R^2^ to assess how well the reconstruction recovers spot and gene patterns. As shown in Table 2, CTFacTomo consistently outperformed both IPF and Tomographer in 3D reconstruction with the lowest MSE and MAE and the highest spot-wise and gene-wise R^2^ on all three datasets. Note that IPF does not work in the high sparsity of the Drosophila embryo data, and thus is not applicable in the comparison. More detailed comparisons of the performance are also shown by scatter plots of R^2^ in Fig. S1, S2 and S3 for the three datasets.

**Table 2.** Reconstruction performance on 3D-projected spatial transcriptomics data. The comparison of IPF, CTFacTomo, CTFacTomo without graph Laplacian regularization, and Tomographer on human heart, mouse brain, and drosophila embryo data. The best mean square error (MSE), mean absolute error (MAE), spot- and gene-wise R^2^ are bold.

| Dataset | Method | Evaluation Metrics |  |  |  |
| --- | --- | --- | --- | --- | --- |
| | | $MSE \downarrow$ | $MAE \downarrow$ | $spot-wise R^2 \uparrow$ | $gene-wise R^2 \uparrow$ |
| **Human Heart**<br>(14,726 × 20 × 20 × 9) | IPF | 0.160 ± 0.081 | 0.190 ± 0.061 | 0.285 ± 0.205 | 0.061 ± 0.132 |
|  | Tomographer | 0.158 ± 0.089 | 0.170 ± 0.056 | 0.337 ± 0.164 | 0.071 ± 0.160 |
|  | CTFacTomo w/o graph reg | 0.176 ± 0.163 | 0.217 ± 0.192 | 0.269 ± 0.230 | 0.015 ± 0.245 |
|  | CTFacTomo w/ graph reg | **0.147 ± 0.093** | **0.156 ± 0.055** | **0.386 ± 0.181** | **0.139 ± 0.224** |
| **Mouse Brain**<br>(10,239 × 30 × 30 × 30) | IPF | 0.426 ± 0.159 | 0.389 ± 0.081 | 0.262 ± 0.245 | 0.066 ± 0.293 |
|  | Tomographer | 0.293 ± 0.136 | 0.363 ± 0.080 | 0.472 ± 0.183 | 0.054 ± 0.207 |
|  | CTFacTomo w/o graph reg | 0.288 ± 0.196 | 0.321 ± 0.104 | 0.517 ± 0.289 | 0.096 ± 0.193 |
|  | CTFacTomo w/ graph reg | **0.245 ± 0.172** | **0.298 ± 0.097** | **0.600 ± 0.174** | **0.171 ± 0.119** |
| **Drosophila Embryo**<br>(14,270 × 63 × 53 × 21) | IPF | - | - | - | - |
|  | Tomographer | 0.017 ± 0.227 | 0.020 ± 0.130 | 0.173 ± 0.187 | 0.097 ± 0.087 |
|  | CTFacTomo w/o graph reg | 0.019 ± 0.245 | 0.021 ± 0.137 | 0.075 ± 0.009 | 0.068 ± 0.085 |
|  | CTFacTomo w/ graph reg | **0.010 ± 0.140** | **0.017 ± 0.100** | **0.449 ± 0.096** | **0.169 ± 0.070** |

**Fig. 2.**
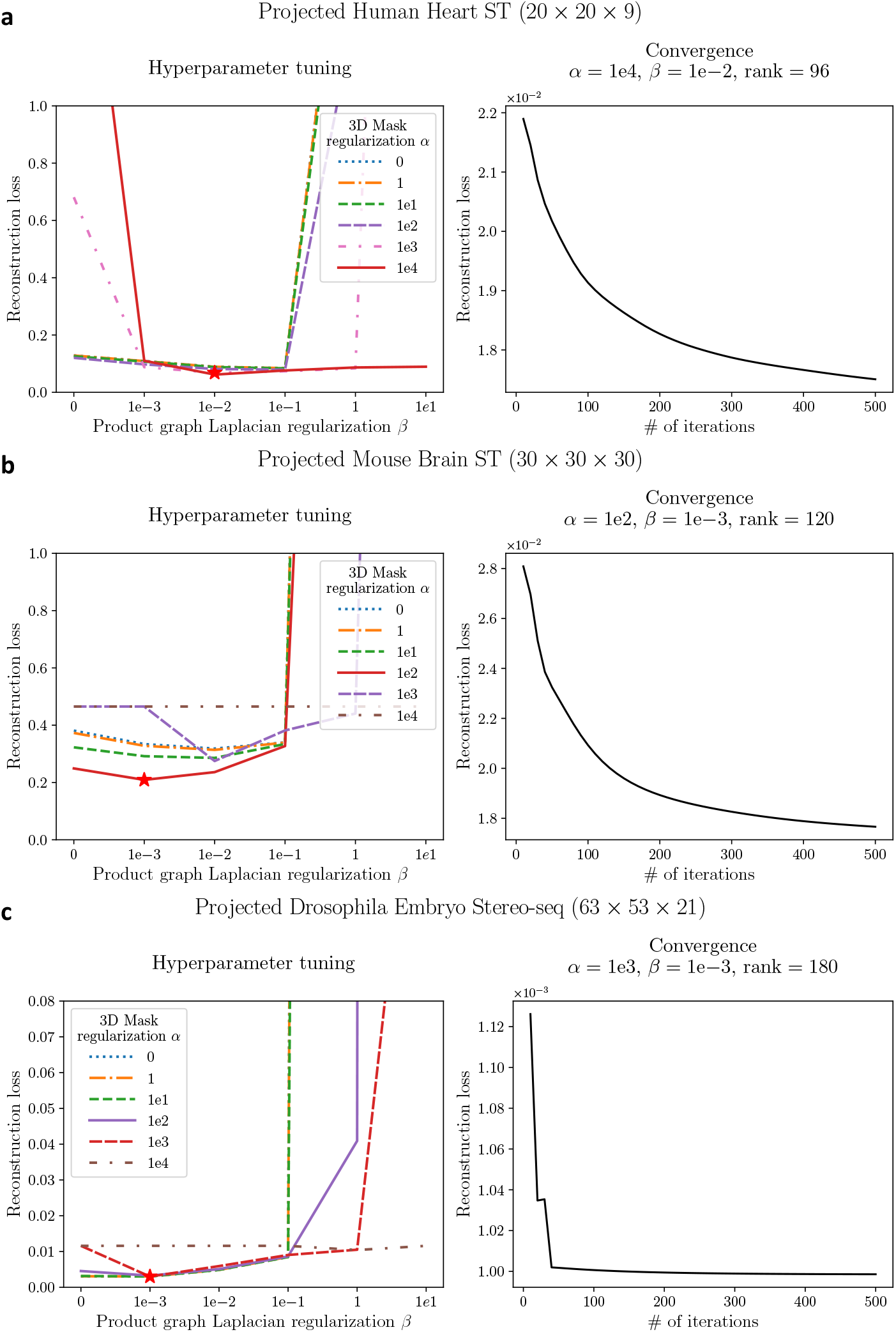
Convergence and Hyperparameter Tuning of CTFacTomo on Projected 3D Spatial Transcriptomics Data. **a** Mouse Brain **b** Human Heart **c** Drosophila Embryo. ***left*** Hyperparameters *α* and *β* are searched over the grid of {0, 1, 1e1, 1e2, 1e3, 1e4} and {0, 1e −3, 1e −2, 1e− 1, 1, 1e−1}; ***right*** Reconstruction loss of CTFacTomo with optimal hyperparameters decreases over the iterations. The optimal hyperparameters (marked by red stars) are determined by the combination that minimizes the reconstruction loss while restricting the expression outside the mask.

To further investigate the contribution of spatial and functional information in the reconstruction, we performed an ablation study by removing the corresponding graph Laplacian regularization in the total loss function. We observed that CTFacTomo outperformed its counterpart without graph Laplacian regularization, which suggests that both spatial and functional relations are crucial guidance for the reconstruction from 1D data. In particular, in the human heart data and the Drosophila embryo data, CTFacTomo without graph regularization performed worse than IPF and Tomographer, suggesting that regularization is often necessary and required for high-order modeling used in CTFacTomo. Note that while Tomographer appears to perform better than IPF with the two-angle (0 degrees and 90 degrees) setting, STRP-seq requires dissecting tissues along the z-axis first and then, further sectioning each slice into parallel strips along either the x-axis (90 degrees) or y-axis (0 degrees), which makes tissue preparation for STRP-seq significantly more complex compared to Tomo-seq in practice.

### 2.2 CTFacTomo reconstructs 3D spatial expressions with consistent patterns from RNA tomography data

In this experiment, we evaluated 3D expression reconstruction on real RNA tomography data. The first dataset contains (50, 49, 56) slices along three spatial axes from a zebrafish embryo at the shield stage Junker et al. (2014). The second dataset contains (56, 54, 60) slices in the three views from a mouse olfactory mucosa Segura et al. (2022).

#### 2.2.1 3D transcriptome reconstruction from RNA tomography for zebrafish embryo

For the RNA tomography data of zebrafish embryo, the hyperparameters *α* = 1, *β* = 1, rank = 500 were chosen for this dataset based on the hyperparameter tuning results shown in Fig. 3a. We evaluated the reconstruction performance by visualizing the expression of several well-known marker genes and comparing their 2D projections with existing ISH images from the same perspective Junker et al. (2014). ISH images are available for four genes, *GSC, SULF1, ISM1*, and *MAGI1B*, and their reconstruction are shown in Figure 4**A**. All four genes are expressed as a patch or different gradients in the organism on the dorsal side at the early development stages of the zebrafish embryo. *GSC* is an important gene in developing mesoderm. While the 3D reconstruction of both CTFacTomo and IPF in some projections, such as along the frontal and anteroposterior axes, highlight the correctly expressed regions in the ISH images, CTFacTomo reconstructions align better with the ISH images without overspreading expression. Moreover, it is obvious that the projections of 3D reconstructions from CTFacTomo along the ventrodorsal axis demonstrated significantly higher agreement with their ISH images compared to IPF.

**Fig. 3.**
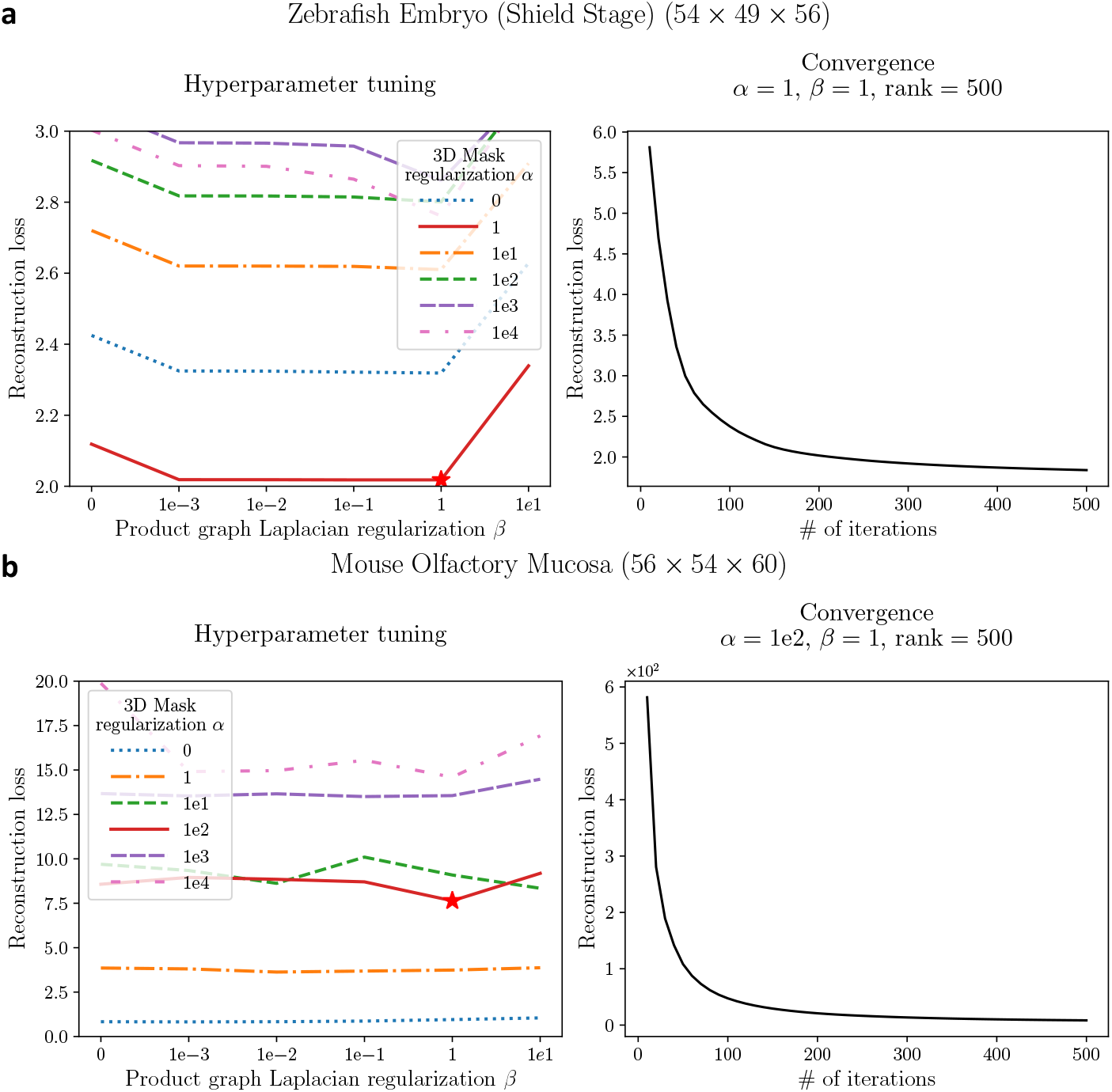
Convergence and Hyperparameter Tuning of CTFacTomo on RNA Tomography Data. **a** Zebrafish Embryo; and **b** Mouse Olfactory Mucosa. ***left*** Hyperparameters *α* and *β* are searched over the grid of {0, 1, 1e1, 1e2, 1e3, 1e4} and {0, 1e−3, 1e −2, 1e −1, 1, 1e1}; ***right*** Reconstruction loss of CTFacTomo with optimial hyperparameters decreases over the iterations. The optimal hyperparameters are determined by the combination that minimizes the reconstruction loss while restricting the expression outside the tissue mask to be less than threshold 1e− 4. Note that in the experiment on mouse olfactory mucosa data, *α* needs to be at least 1e2 to restrict the level of expression outside the tissue mask to be less than threshold 1e− 4. Thus, *α* =1e2 was selected as the best parameter in the tuning.

**Fig. 4.**
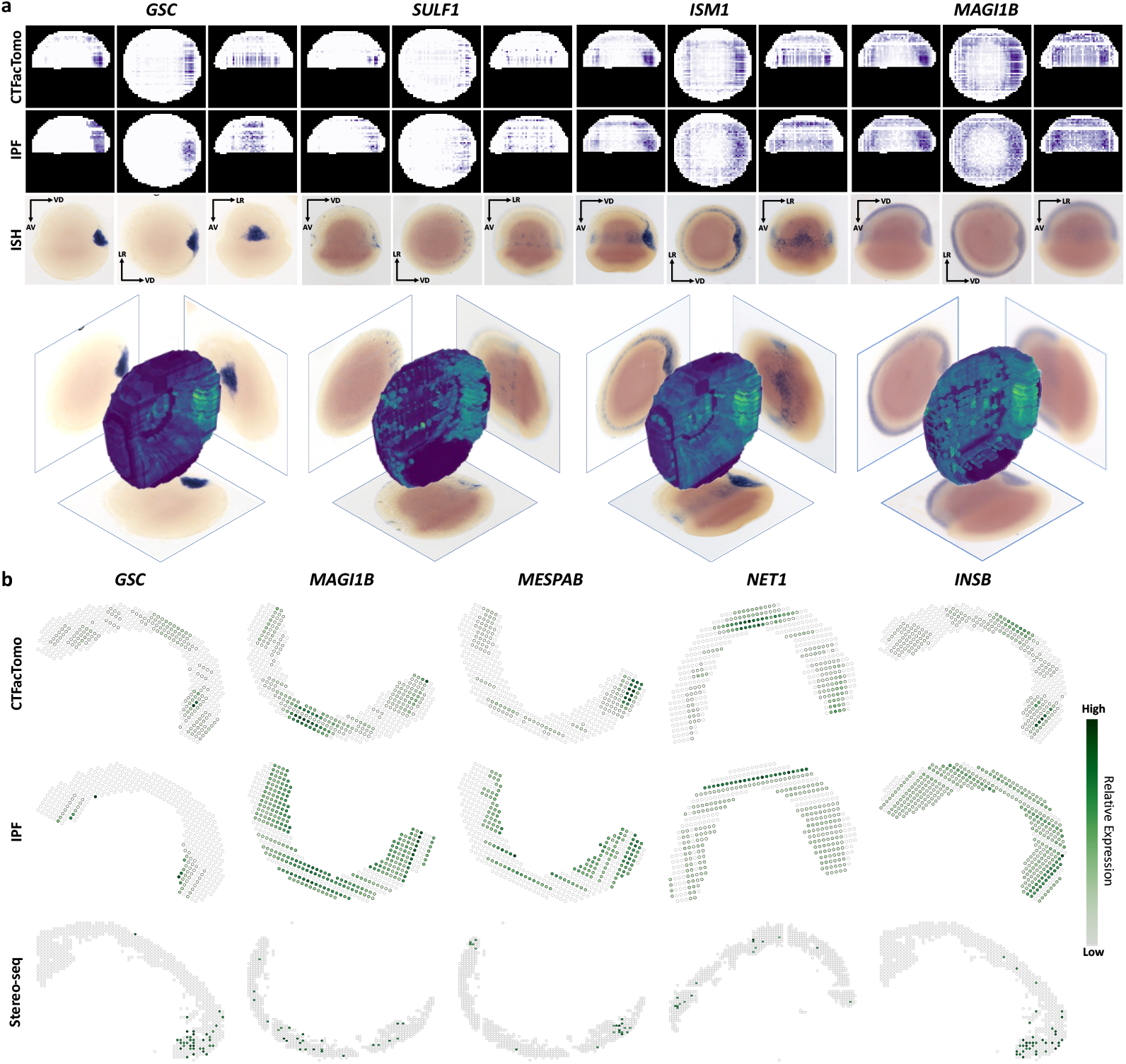
Visualization and evaluation of marker genes in zebrafish Tomo-seq data. **a** 2D projection of the reconstructed expression for the entire zebrafish embryo at shield stage along frontal, anteroposterior, and ventrodorsal axes of four genes (*GSC, SULF1, ISM1*, and *MAGI1B*). Shaded blue areas in the ISH images denote expression regions. The bottom row shows the 3D visualization of the reconstruction by CTFacTomo. **b** A comparison between the reconstructed slices by CTFacTomo and IPF with the best matched Stereo-seq slices by Moran scores of five marker genes (*GSC, MAGI1B, MESPAB, NET1, INSB*). Spots with a black outer edge represent non-zero expression *>* 0.0001 after normalization, and the gradient represents relative expression from low (light gray) to high (dark green).

To further compare the reconstructions of CTFacTomo and IPF from RNA tomography data, we did both a quantitative and qualitative slice-wise comparison of the reconstructions of CTFTacTomo and IPF with slices of the external Stereo-seq dataset Liu et al. (2022). This provides a more granular evaluation of the reconstruction as compared to when looking at the 2D projections. All the slices for comparison are along the sagittal plane. Note, the original Tomo-seq slices have a thickness of 18*µm* and the Stereo-seq slices have a thickness of 12*µm*. Quantitatively, we used a bivariate variation of Moran’s I score as a measure of spatial correlation to compare a pair of reconstructed and Stereo-seq slices (see **Methods** section). Specifically, we took the twelve middle slices of the reconstructed zebrafish embryo and compared them to the twelve available slices of Stereo-seq data that were obtained from the matched middle sections of the embryo. For each gene, we obtained 144 Moran scores, one for each of the pairs. We then consider the maximum score among these 144 Moran scores shown in Figure S4, and picked matching reconstructed and Stereo-seq slices accordingly to visualize as well as use these maximum scores for a global comparison between CTFacTomo and IPF. In Figure 4b, we visualize the pairs with the best Moran score for five marker genes (*GSC, MAGI1B, MESPAB, NET1, INSB*). We also report the detailed Moran scores in Table 3. These comparisons again confirm the better matching of the reconstruction by CTFacTomo than IPF. In Fig. 5, we also show a global analysis of the Moran scores for all 5,439 genes matched between Tomo-seq data and Stereo-seq data. The results show that CTFactTomo reconstructed significantly more consistent spatial gene expressions than IPF in the top 2,000 highly variable genes and all 5,439 genes.

**Table 3.** Moran scores between the reconstructed data and the matched Stereo-seq data for five marker genes. All spots or top spots are used in the calculation.

| Marker Gene | All spots |  | Top spots |  |
| --- | --- | --- | --- | --- |
|  | IPF | CTFacTomo | IPF | CTFacTomo |
| *GSC* | 0.0644 | **0.2350** | 0.4621 | **2.3425** |
| *MAGI1B* | 0.0619 | **0.1528** | 0.3596 | **1.0123** |
| *MESPAB* | 0.1870 | **0.3022** | 1.3065 | **2.9081** |
| *NET1* | 0.0473 | **0.0966** | 0.6298 | **1.2809** |
| *INSB* | 0.0744 | **0.0885** | 0.4669 | **0.7230** |

**Fig. 5.**
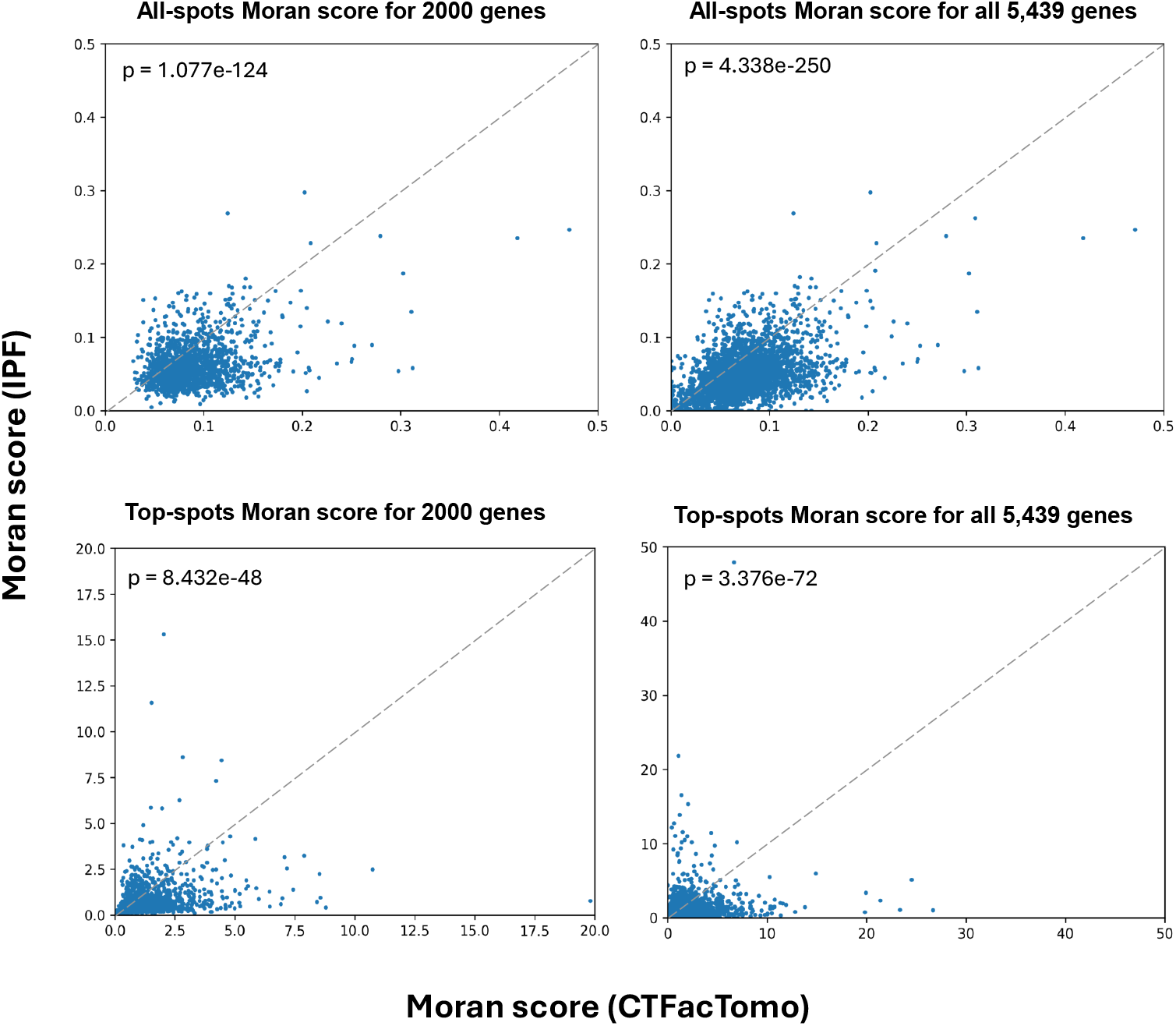
Comparison of Spatial Coherence between Reconstructed and Stereo-seq slices by Moran scores. The scatter plots compare the Moran scores of the top 2,000 highly variable genes or all 5,439 genes when comparing the reconstructed data by CTFacTomo and IPF to Stereoseq slice-wise data. The first row shows the Moran scores calculated over all spots and the second row shows the Moran scores calculated over the top spots.

#### 2.2.2 3D Transcriptome reconstruction from RNA tomography for mouse olfactory mucosa

In the experiment on the RNA tomography dataset of mouse olfactory mucosa, the hyperparameters *α* = 100, *β* = 1, rank = 500 were selected based on the hyperparameter tuning results shown in Fig. 3b. The stopping criteria was to have a residual threshold of 1e− 4 and a max epoch of 1000. Similar to the previous experiment, we visualized the reconstructions from CTFacTomo (9, 254× 56 ×54 ×60) and IPF for comparisons on five marker genes with ISH images available Segura et al. (2022). The five marker genes (*OLFR309, OLFR618, OLFR727, CYTL1, MOXD2*) are shown in Fig. 6. *OLFR309, OLFR727*, and *OLFR618* are three olfactory receptors and genetic markers of different mature olfactory sensory neuron subtypes. *CYTL1* is a cytokine-like protein playing roles in osteogenesis, chondrogenesis, and bone and cartilage homeostasis Shin et al. (2019). *MOXD2* is a mono-oxygenase dopamine hydroxylase-like protein functioning in olfaction Kim et al. (2014).

**Fig. 6.**
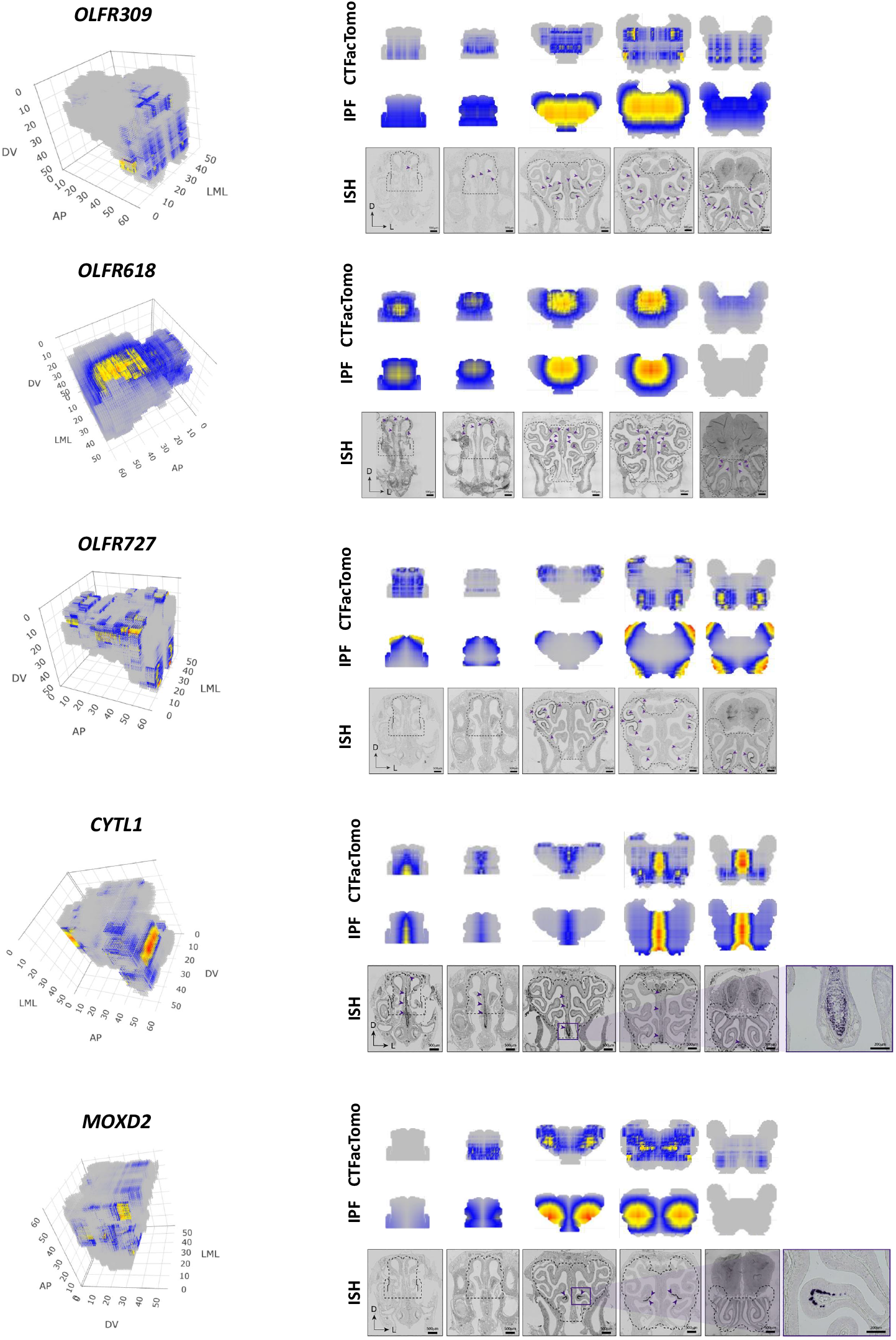
Visualization and Evaluation of Marker Genes in Mouse Olfactory Mucosa. We visualize both the full 3D reconstruction as well as slice-wise visualizations for five marker genes (*OLFR309, OLFR618, OLFR727, CYTL1, MOXD2*). ***left*** The 3D gene expression output by CTFacTomo for the five marker genes. ***right*** A comparison between CTFacTomo and IPF in a slice-wise view of the 3D reconstruction in comparison to the ground-truth ISH images for each gene. Within the ISH images, purple triangles represent regions of expression.

It is most clear when comparing a reconstructed slice of *OLFR618* along the anteroposterior axis with its last ISH image, where CTFacTomo accurately reconstructs expression, whereas IPF fails to reconstruct any. The fourth image of *OLFR727* shows another example of this, where IPF exaggerates the expression in the bottom two corners and CTFacTomo more accurately shows the two concentrated regions of expression. We see a similar example in the fifth image of *OLFR727*, where IPF even incorrectly infers expression in the upper region, whereas CTFacTomo correctly infers that there is only expression in the bottom region of that slice. However, there are some instances among the visualized genes in which both CTFacTomo and IPF incorrectly represent spatial gene expression. For example, both CTFacTomo and IPF struggle to accurately reconstruct the expression of *Olfr309*. Note that since *OLFR618* ‘s expression level is only around 1% of the other genes, we set *α* = 1e −3 to run CTFacTomo separately such that the extremely low expressions can be retained in the reconstruction. The expression of all the slices are also visualized in Fig. S5 for reconstructions by CTFacTomo and Fig. S6 for reconstructions by IPF. In summary, through a visual inspection of the reconstructions, we see that CTFacTomo can find more focused expression regions than IPF, likely due to IPF’s tendency to overspread the regions on such data.

### 2.3 Reconstructed 3D transcriptomes from CTFacTomo enhances biological interpretation of spatially co-expressed gene clusters

To evaluate whether the reconstructed 3D spatial expressions can enhance biological interpretation, we further investigated the reconstruction on real RNA tomography data. We first applied kMeans to group genes into 100 co-expressed clusters with either concatenated 1D expressions in the original RNA tomography profiles or reconstructed 3D expressions. Gene Ontology (GO) enrichment analysis was performed by using the enrichGO function from R package clusterProfiler for each cluster, with significance determined by the p-value of the most significant GO term of each cluster. The comparison of the number of significant clusters by different reconstruction methods is shown in Fig. 7. We observed that the 3D expressions reconstructed by CTFacTomo consistently identified more significant co-expressed gene clusters across all p-value thresholds for both zebrafish embryo and mouse olfactory mucosa than the data reconstructed by IPF and the original RNA tomography profiles, which suggests CTFacTomo could improve the biological interpretation with more functional enrichment. Additionally, we introduced another comparison by replacing the PPI network in the product graph Laplacian regularization of CTFacTomo with an identity matrix of the same size. The results showed that CTFacTomo without the PPI network detected fewer significant gene clusters than CTFacTomo due to the loss of functional information in the PPI network. Nevertheless, this variation of CTFacTomo still detects overall more functionally enriched clusters than IPF in both datasets, and the original Tomo-seq profiles in mouse olfactory mucosa data.

**Fig. 7.**
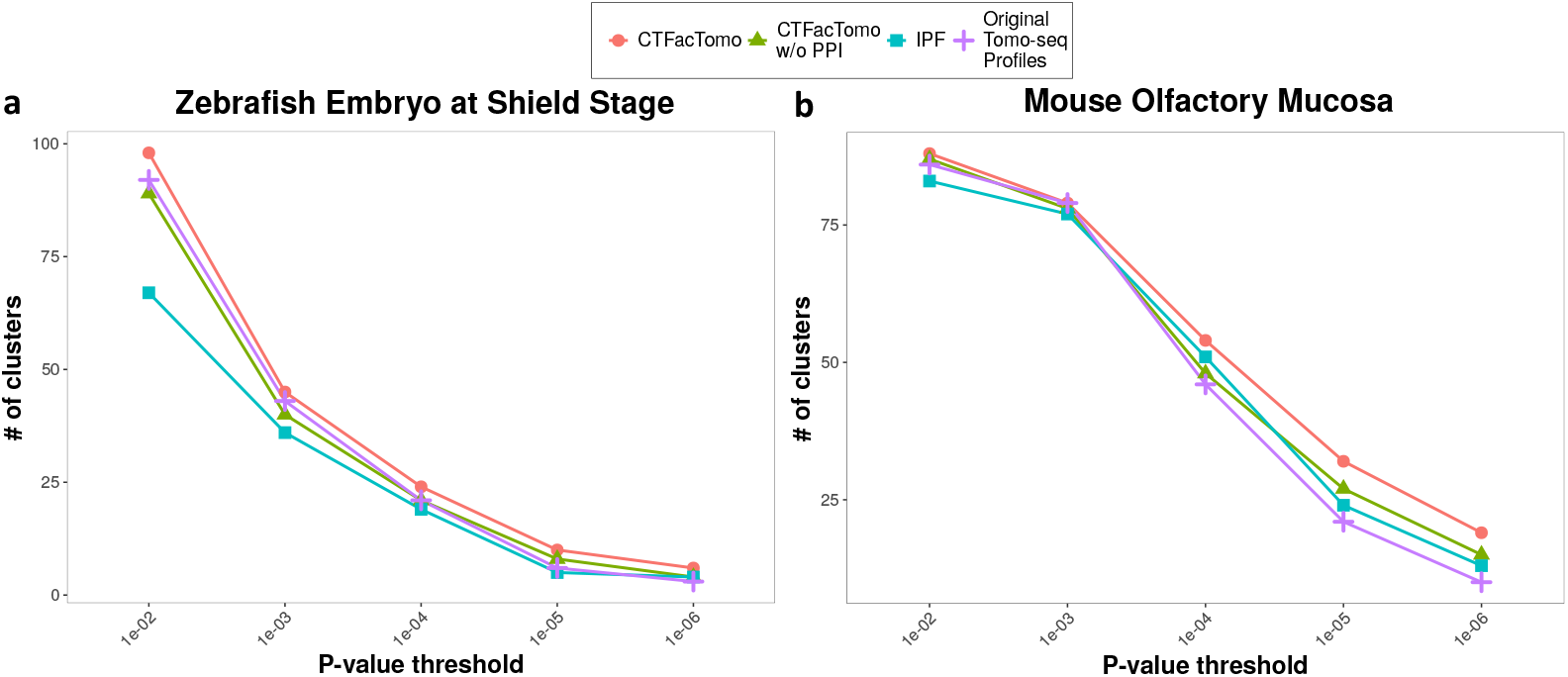
Comparison of Significantly Enriched Gene Clusters Identified Using Reconstructed 3D Expressions. The number of significantly enriched gene clusters at varying p-value thresholds in **a** Zebrafish Embryo and **b** Mouse Olfactory Mucosa.

## 3 Discussion

Recently, microdissection-based methods such as variations of Tomo-seq, e.g., Geoseq (Peng et al. 2016) and STRP-seq (Schede et al. 2021) have gained increasing attention. These newer technologies leverage adjacent slices as replicates such that no identical biological replicate is necessary for tomography. Furthermore, more flexible tomography views can be adopted beyond the three orthogonal views (Schede et al. 2021). For instance, Geo-seq has been applied to explore temporal and spatial patterns in various embryo studies (Peng et al. 2019; Xue et al. 2019; Wang et al. 2023; Qu et al. 2023), and STRP-seq has demonstrated its potential in uncovering intricate spatial patterns in a non-model organism (Schede et al. 2021). These new developments greatly improved the applicability of microdissection-based methods. Additionally, high-throughput readout and relatively lower cost make these methods highly favorable for some specific research applications in developmental biology.

As demonstrated in the analysis and the experiments in this study, CTFacTomo performed better than IPF and Tomographer for 3D reconstruction. There are several advantages of modeling with CTFacTomo. First, CTFacTomo utilizes crucial spatial information and gene functional relations to better solve the difficult reconstruction problem from under-represented 1D expressions from RNA tomography. Second, CTFacTomo is a multitask model for whole transcriptome reconstruction in 3D, while single-gene construction by IPF is very prone to noise. Finally, CTFacTomo is a high-order method that better characterizes the 3D organization of the information, while 2D reconstruction by Tomographer loses the important 3D dependence.

It is also important to note that CTFacTomo is based on a different formulation in contrast to the standard formulation of tensor factorization, where standard tensor factorization is derived from a given tensor with all (or some) known entries, while the formulation of CTFacTomo learns a factorization from the aggregated sums in certain views from collapsing the tensor without knowing any actual entry in the tensor to be constructed. Thus, the algorithm for such formulation is entirely different from standard tensor factorization methods and has a great potential to generalize to different kinds of tomography constructions in future work.

## 4 Methods

In CTFacTomo, we model the reconstructed 3D spatial gene expression as 4-way tensor 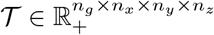, and we also model the 1D spatial gene expressions along different spatial axes as 2D matrices 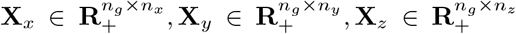, and the tensor-based model for 3D spatial gene expression reconstruction is illustrated in the Figure 1. The key modeling ideas are: A) the collapse of the reconstructed 3D spatial gene expression tensor *T* along the gene and a given spatial axes collapse(𝒯, *g, i*) should be identical to the 1D gene expression matrix **X**_*i*_; B) the reconstructed tensor 𝒯 should be approximated by 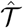 that can be compressed in a rank-*R* CP decomposition form 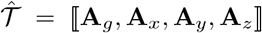 for time and space efficiencies; C) the entries in the reconstructed tensor 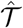 should align the spatial relations in the 3D space and functional relations among genes.

### 4.1 Preliminaries

Here, we only present the minimal necessary definitions for introducing CTFacTomo.

#### Canonical Polyadic Decomposition (CPD)

An *M* -way tensor 𝒯 can be approximated in a compressed rank-*R* CPD representation as follows,

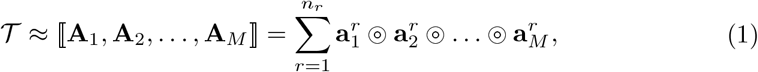

where ⟦ · ⟧denotes the Kruskal operator, ⊚ denotes vector outer product, *n*_*r*_ is the rank of the decomposition, 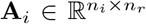 represents the factor matrix along mode-*i*, and **a**^*r*^ is a vector in the *r*-th column of **A**_*m*_.

#### Tensor Matricization

A tensor represented by CP decomposition can be matricized along *i*-th mode as 𝒯 _(*i*)_ = **A**_*i*_(**A**_1_ ⊙ … ⊙ **A**_*i−*1_ ⊙ **A**_*i*+1_ ⊙ … ⊙ **A**_*M*_)^*T*^, where denotes Khatri-Rao product, and 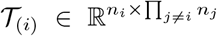. As a special case of matricization, a tensor in the form of CP decomposition can be vectorized as vec(𝒯) = (**A**_1_ ⊙ … ⊙ **A**_*j*_ ⊙ … ⊙ **A**_*M*_)**1**^*T*^, where vec(·) represents the vectorization function and **1** is an all-ones vector of size *n*_*r*_.

#### Tensor Collapsing

Collapsing operation collapses a tensor into a lower-order one with given modes by summing over entries along the remaining modes. Collapsing any one mode on the CPD of a tensor can be written as a collapsing function collapse(·) as follows,

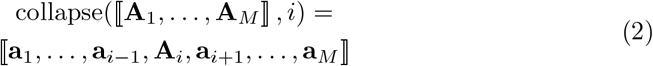

where 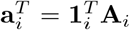 denotes the row summation of **A**_*i*_. Similarly, collapsing given any two modes on the CPD of a tensor can be written as,

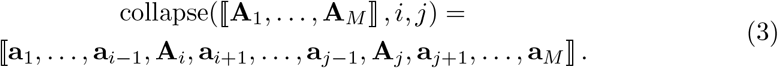

Accordingly, the collapsing operation can be defined on the tensor of CP decomposition form given more than two modes.

#### Product Graph

Given *M* undirected graphs *{G*_*m*_ = (*V*_*m*_, *E*_*m*_): *m* = 1, 2, …, *M}*, where *V*_*m*_ and *E*_*m*_ denotes the set of nodes and edges in the graph *G*_*m*_, and |*V*_*m*_| = *n*_*m*_ indicates the number of nodes in the *G*_*m*_. Let *G*_*p*_ be a new graph combining the set of graphs *{G*_*m*_: *m* = 1, 2, …, *M}*, denoted as product graph *G*_*p*_ = (*V*_*p*_, *E*_*p*_) with the number of nodes 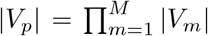. For any pair of nodes (*a*_1_, *a*_2_, …, *a*_*M*_) and (*b*_1_, *b*_2_, …, *b*_*M*_) in the product graph *G*_*p*_, the corresponding edge is determined by the adjacency or equality of *a*_*m*_ and *b*_*m*_ in the graph *G*_*m*_. For simplicity, we only used Cartesian product graph in this work, where ((*a*_1_, *a*_2_, …, *a*_*M*_), (*b*_1_, *b*_2_, …, *b*_*M*_)) ∈ *E*_*p*_ if and only if (*a*_*m*_, *b*_*m*_) ∈ *E*_*m*_ while *a*_*k*_ = *b*_*k*_, ∀*k* ≠*m*.

Let **W**_*m*_ be the adjacency matrix of *G*_*m*_, where [**W**_*m*_]_*ij*_ = 1 if there is an edge between *i*- and *j*-th nodes in *G*_*m*_ and 0 otherwise, and let 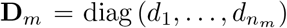 be the degree matrix of *G*_*m*_ with *d*_*ij*_ = _*j*_ [**W**_*m*_]_*ij*_. **L**_*m*_ = **D**_*m*_ − **W**_*m*_ represents the graph Laplacian for *G*_*m*_. Then the adjacency matrix and Laplacian matrix of Cartesian product graph *G*_*p*_ can be calculated by 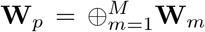 and 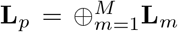respectively, where ⊕ denotes the Kronecker sum.

#### Product Graph Laplacian Regularization

Suppose a tensor *M* -way 𝒯 with each mode *m* associated with a graphs *G*_*m*_ encoding the prior knowledge. Let *G*_*p*_ be the product of the graphs from different modes, where **W**_*p*_ and **L**_*p*_ are adjacency matrix and Laplacian matrix of *G*_*p*_. Each entry in 𝒯 corresponds to one node in *G*_*p*_, and then graph Laplacian regularization can be used to smooth values in 𝒯 over the manifolds of the graph *G*_*p*_, ensuring that adjacent nodes in this high-order graph share similar values, which can be mathematically defined in the quadratic form as vec(𝒯)^*T*^ **L**_*p*_vec(𝒯) = _*ij*_ [**W**_*p*_]_*ij*_([vec(𝒯)]_*i*_ − [vec(𝒯)]_*j*_). When 𝒯 is represented using the form of CP decomposition, product graph Laplacian regularization can be formulated as follows:

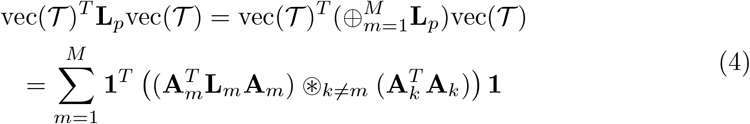

where ⊛ denotes Hadamard product, and **1** is an all-ones vector of size rank *n*_*r*_.

### 4.2 Loss function

The overview of CTFacTomo is illustrated in Fig. 1. The inputs of CTFacTomo are the tomography transcriptomics data along different spatial axes as gene expression matrices 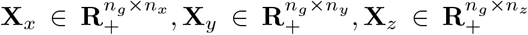, and a 3D image mask as a 3-way tensor 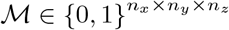 as shown in Fig. 1a. The 1D spatial chain relations along three orthogonal axes and functional relations among genes are given in four knowledge graphs *{G*_*i*_, *i* = *g, x, y, z}* (Fig. 1c). CTFacTomo reconstructs 3D spatial gene expression in a 4-way tensor 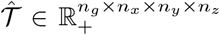 represented by CP decomposition while being guided by prior knowledge encoded in the spatial and functional graphs (Fig. 1b). We next formulate the learning problem and define each term in the loss function of CTFacTomo.

#### Reconstruction Loss

We first define reconstruction loss 𝒥_1_, which ensures the collapse of the reconstructed 3D spatial gene expression tensor 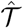 along the gene and a given spatial axes should be identical to the 1D gene expression matrix **X**_*i*_, and the loss term can be written as,

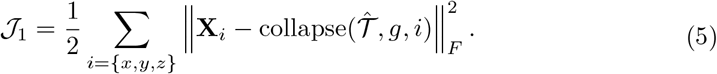

Here, 4-way tensor 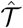 is collapsed (summed) over the other two spatial modes along each of the x-axis, y-axis and z-axis, exactly modeling the tomography projection onto the cryosections in each view.

#### 3D Masking Loss

To prevent the reconstructed 3D spatial gene expression from over-spreading to regions outside of the tissue, a loss term 𝒥_2_ is introduced to fit the given 3D mask ℳ that outlines the tissue shape as follows,

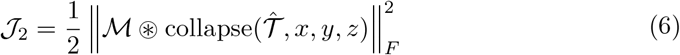

where ⊛ denotes Hadamard product. Here, the 4-way tensor 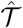 is collapsed (summed) over all the genes in each spatial location (spot), and the loss penalizes the non-zero sums in the spots outside of the tissue.

#### Product Graph Laplacian Regularization

To leverage both spatial relations in 3D space and functional relations among genes for reconstruction, a Laplacian regularization of the Cartesian product of gene and spatial graphs *{G*_*i*_, *i* = *g, x, y, z}* in 𝒥_3_ is used to smooth the entries in 𝒯 over the manifolds in the Cartesian product graph *G*_*p*_, ensuring that the expressions of adjacent nodes in this high-order graph share similar values, defined as follows,

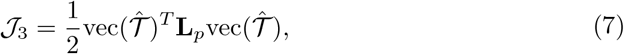

Here **L**_*p*_ = **L**_*g*_ ⊕ **L**_*x*_ ⊕ **L**_*y*_ ⊕ **L**_*z*_ is the graph Laplacian of *G*_*p*_, where **L**_*i*_, ∀*i* = *g, x, y, z*, is the Laplacian graph of *G*_*i*_, and ⊕ denotes Kronecker sum (Li et al. 2021).

#### Total Loss function

Lastly, the total loss function 𝒥 to learn a rank-*R* CP decomposition 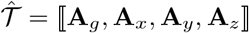 is the sum of 𝒥_1_, 𝒥_2_ and 𝒥_3_ as follows,

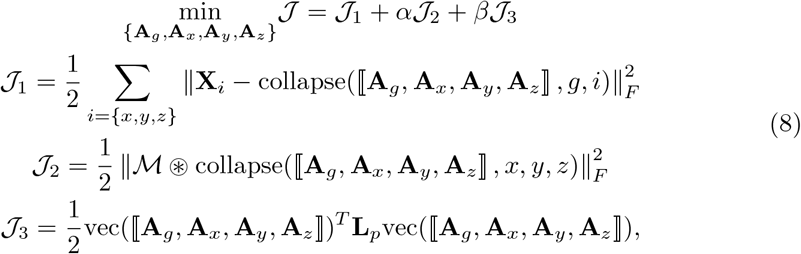

where *α* and *β* are hyperparameters weighting the loss terms in the loss function.

### 4.3 Multiplicative updating

To minimize the loss function in Eq. 8, we propose an iterative optimization method based on multiplicative updating rules (Lee and Seung 2000; Cai et al. 2008; Li et al. 2019), which ensures a stationary solution of 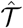 by alternatively updating each {**A**_*i*_, *i* = *g, x, y, z}* with the derivatives of 𝒥 with respect to the **A**_*i*_.

For simplicity, we decompose the loss function into multiple terms and introduce their derivatives with respect to **A**_*g*_ and **A**_*i*_, ∀*i* = *x, y, z* individually. To facilitate the calculation of the derivatives, we define the auxiliary variables in Table 4.

**Table 4.**
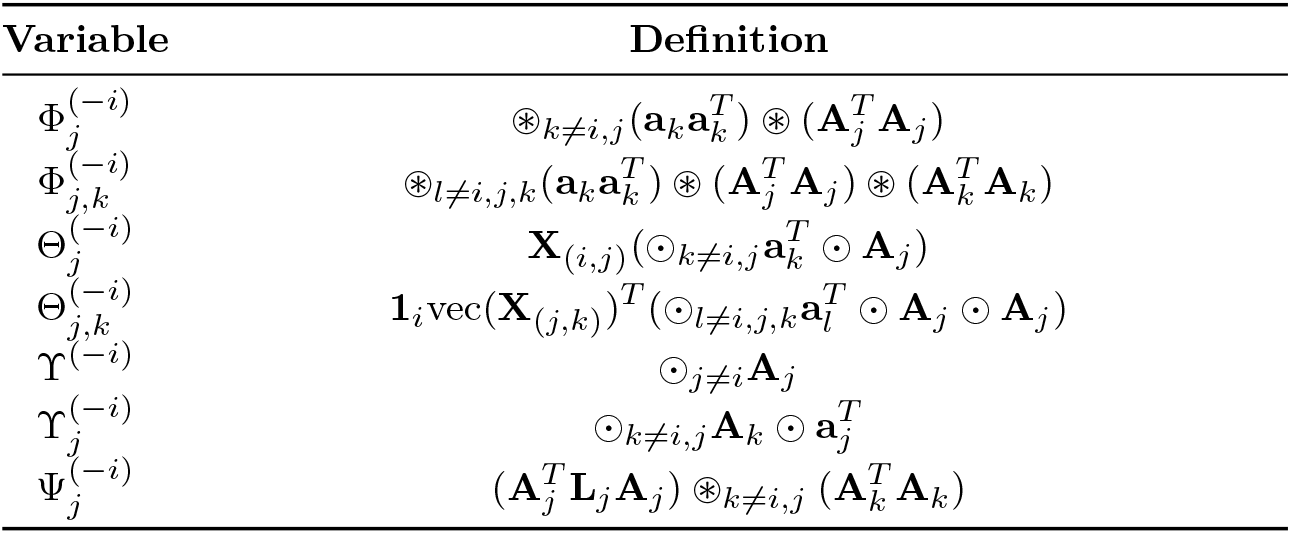
Auxiliary variables.

With the definitions in Eq. 4, the partial derivatives of 𝒥_1_ with respect to **A**_*g*_ and **A**_*i*_, ∀*i* = *x, y, z* can be represented as follows:

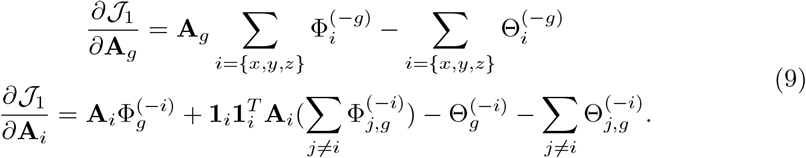

Similarly, the partial derivatives of 𝒥_2_ with respect to **A**_*g*_ and **A**_*i*_, ∀*i* = *x, y, z* can be represented as follows:

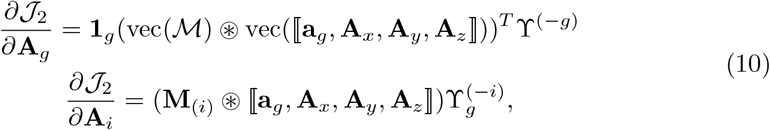

where 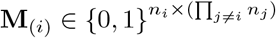 denotes the mode-*i* matricization of tensor ℳ.

Following the derivation in (Li et al. 2019), the partial derivatives of *J*_3_ with respect to **A**_*i*_, ∀*i* = *g, x, y, z* can be represented as follows:

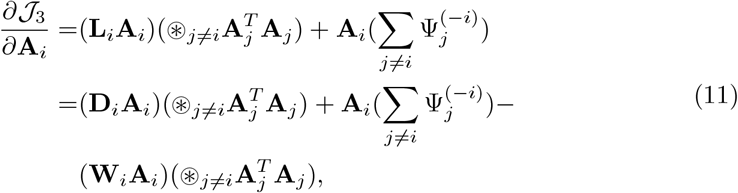

where **W**_*i*_ and **D**_*i*_ are the adjacency matrix and degree matrix of the graph *G*_*i*_, and **L**_*i*_ = **D**_*i*_ − **W**_*i*_.

After combining the derivatives in Eq. 9, 10, 11, the derivative of 𝒥 with respect to **A**_*i*_, ∀*i* = *g, x, y, z* can be formalized as follows:

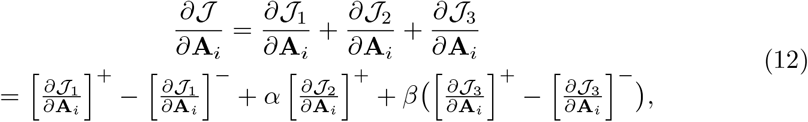

where 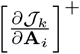 and 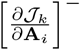 are the non-negative components in the partial derivative of 𝒥_*k*_, ∀*k* = 1, 2, 3 with respect to **A**_*i*_, ∀*i* = *g, x, y, z*.

The loss function 𝒥 will be monotonically decreased by alternatively updating each factor matrix **A**_*i*_, ∀*i* = *g, x, y, z* using the following multiplicative updating rule in each iteration until convergence.

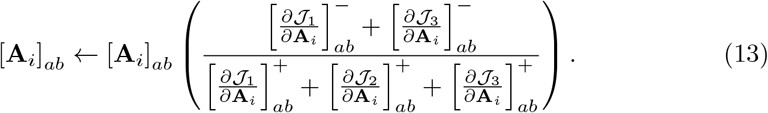

### 4.4 Hyperparameter tuning

To tune the hyperparameters *α* and *β* in the loss function (Eq. 8), we applied grid search over the ranges {0, 1, 1e1, 1e2, 1e3, 1e4} and {0, 1e−3, 1e−2, 1e−1, 1, 1e1} to find the best combination that minimizes the reconstruction loss 𝒥_1_ while restricting the over-expression outside tissue mask to be less than a very small threshold such as 1e−4. This restriction can be achieved by imposing a relatively large *α* such that a small 𝒥_2_ loss is assured.

We determined the rank of CPD using min(Π _*i*_ *k*_*i*_, 500), ∀*i* = *x, y, z*, where *k*_*i*_ denotes the number of principal components explaining more than 95% variance of the RNA tomography data along different spatial axes **X**_*i*_, *i*∀ = *x, y, z*. By default, CTFacTomo terminates when the sum of the differences between the previous and current components in each mode is smaller than a small threshold such as 1e −3 or 1e− 4, or when the maximum iteration limit (500 or 1000) was reached. In both ST data and RNA tomography data, IPF terminates either when the differences between the given and the reconstructed 1D spatial expressions in the sum over the three spatial axes are all less than a threshold 1 or upon reaching the maximum iteration limit 200. Tomographer was tuned by following the guidelines provided in the tutorial of the package, and the default settings were applied in all the experiments.

### 4.5 Evaluation metrics

#### Reconstruction Errors

To assess the reconstruction performance of projected 3D spatial transcriptome, we applied three widely used metrics including mean square error (MSE), mean absolute error (MAE), and coefficient of determination R^2^ over spots or genes. These metrics are defined as follows,

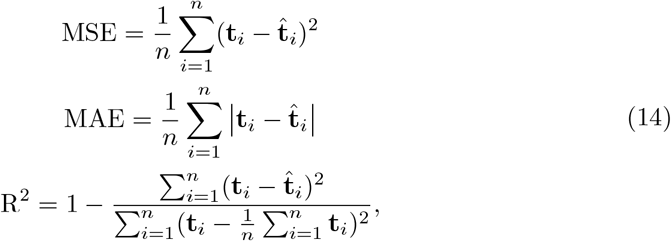

where **t** ∈ ℝ^*n*^ denotes the expression of each spot (*n* = *n*_*g*_) or gene (*n* = *n*_*x*_ × *n*_*y*_) in the original raw spatial transcriptomics data 𝒯 while 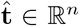 denotes the expression of each spot (*n* = *n*_*g*_) or gene (*n* = *n*_*x*_ × *n*_*y*_) from the imputed spatial transcriptomics data 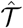 after combining the predictions of each fold in the cross-validation.

#### Spatial Coherence

To compare slices of the CTFacTomo and IPF reconstructions on RNA tomography data with data coming from matched slices of the Stereo-seq data, we used a bivariate version of Moran’s I score Li et al. (2023), of which the univariate version was first introduced in Moran (1950). The bivariate formulation is shown below.

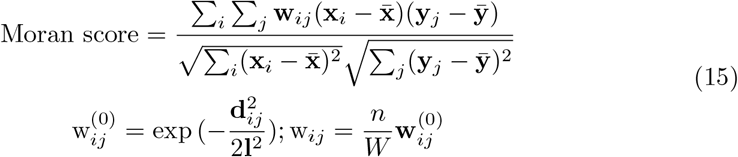

Here, **x**_*i*_ and **y**_*j*_ are the gene expression of a gene at the i-th spot in the reconstructed slice and the expression at the j-th spot in the Stereo-seq slice. Accordingly, 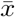 and 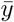 are the mean expressions of the gene in the slice. 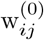 is the unnormalized weight between the spatial location of spot i and spot j. We then normalize using the number of spots *n* and *W*, the sum of the weight matrix. Throughout this paper, *n* refers to the number of Stereo-seq spots which is different for every Stereo-seq slice. One important difference to note as compared to more common previous uses of the bivariate Moran score is that the number of spots indexed by *i* and the number of spots indexed by *j* are different for the use case. This is necessary since the grid of the spots in the Stereo-seq slice and the reconstruction from Tomo-seq data have different arrangements depending on the resolution of the data. In addition, other than calculating the score between all the spots from the compared slices, we also calculated the Moran score only between the top spots. Specifically, we set w_*ij*_ = 0, if **y**_*j*_ = 0 in the Stereo-seq slice or **x**_*i*_ is in the top 2*n*_*>*0_ spots in the reconstructed slice, where *n*_*>*0_ is the number of non-zero spots in **y**_*j*_. This variation allows the score to focus on non-zero (or top) entries for a more sensitive evaluation of the sparse data.

#### Enrichment Significance

To measure the 3D reconstruction performance with Tomo-seq data for gene clustering, we computed the log of the *q*-value of the most significant enriched GO term for each gene cluster and then averaged these minimal q-values across all gene clusters to evaluate the overall enrichment significance. We performed enrichment over 10, 185 GO terms from the C5 collection in the Molecular Signatures Database (MSigDB), which includes 7, 751 biological process (BP) terms, 1, 009 cellular component (CC) terms, and 1, 772 molecular function terms. We calculated q-values by adjusting enrichment p-values by false discovery control (FDR) with the Benjamini-Hochberg (BH) procedure.

### 4.6 Time and Space Complexity

Let *n*_*r*_ = *R* and | · | denote the number of non-zero entries in either a matrix or a tensor, the time complexity to update **A**_*i*_, ∀*i* = *g, x, y, z* in each iteration is 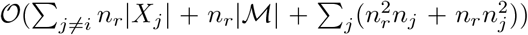, which is derived based on the time complexity of matricized tensor times Khatri-Rao product (MTTKRP) (Choi and Vishwanathan 2014; *Smith et al. 2015). Since in RNA tomography data, the number of genes is larger in several magnitudes, i*.*e. n*_*i*_ *<< n*_*g*_, ∀*i* = *x, y, z, r*, the time complexity is likely to be upper bounded by either 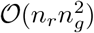 or 𝒪 (*n*_*r*_*n*_*x*_*n*_*y*_*n*_*z*_), in neither of which the complexity in the size of the full 4-ways of the tensor is needed. The space required for the inputs RNA tomography data, spatial and functional graphs, as well as 3D image mask is 𝒪 (∑_*i*≠*g*_ |**X**_*i*_| +| ℳ|+ ∑_*i*_(|**W**| + *n*_*g*_*n*_*r*_)). In practice, CTFacTomo converges within 10 minutes and requires up to 10GB of memory for computation on the real datasets tested in this study.

### 4.7 Data preparation

All the datasets used in this study are summarized in Table 1 along with the experimental specifications, dataset sizes, and platforms.

The human heart ST dataset contains a stack of 9 adjacent slices of 2D ST spatial transcriptomics data from a developing human heart at 6.5 post-conception weeks (PCW) sectioned along the ventrodorsal axis. The mouse brain ST dataset contains a stack of 40 slices sectioned along the anteroposterior axis of an adult mouse brain. In both datasets, each ST slice contains 33 ×35 spots, and these slices were manually registered based on their associated H&E staining images and batch-corrected across slices. (Ortiz et al. 2020; Asp et al. 2019). The Drosophila embryo 3D spatial transcriptomics dataset consists of a stack of 21 slices sectioned along the left-right axis of the Drosophila embryo at the second instar larva (L2) stage (Wang et al. 2022). All three datasets were normalized in log-space.

In the two ST datasets, we binned the spots in the stack of 2D expression data into a 3D tensor to reconcile the coordinate shift across the slices. The spots were assigned to the closest bin and averaged within the bin. The dimensions after binning are 14, 726×20× 20 ×9 for the human heart and 10, 239×30× 30 ×30 for the mouse brain, after removing non-expressed genes and background regions for both tissues. The Drosophila embryo data was directly formed into a 14, 270× 124 ×105 ×21 tensor, with non-expressed genes and background regions being removed, and then binned into a 14, 270×63×53 ×21 tensor to amplify the expression signals in simulation studies. Each slice contains 124 × 105 spots, each of which is a bin of 50× 50 DNA Nanoballs (DNBs) as processed in the original study. The 3D mask was derived for the no-expression spots by collapsing gene expressions along the gene axis in the tensor. A spatial chain graph was constructed by the number of slices in each spatial axis and the protein-protein interaction (PPI) networks for *Homo sapiens, Mus musculus*, and *Drosophila melanogaster* were obtained from BioGRID version 4.4.242 (Oughtred et al. 2021) using mainly only physical interactions.

In the second experiment, we evaluated 3D expression reconstruction on real RNA tomography data. The first dataset contains (50, 49, 56) slices along three spatial axes from a zebrafish embryo at the shield stage (Junker et al. 2014). The second dataset contains (56, 54, 60) slices in the three views from a mouse olfactory mucosa (Segura et al. 2022). We constructed a *Danio rerio* PPI network from STRING (Szklarczyk et al. 2023) with the interactions of confidence scores at least 0.8 retained for sufficient connectivity in the network. The *Mus musculus* PPI network was also constructed from BioGRID (Oughtred et al. 2021). After the intersection of the PPI network and the gene expression profiles, 6, 153 and 9, 254 genes are retained in the zebrafish embryo and the mouse olfactory mucosa, respectively.

In the zebrafish embryo data, we followed the same procedure in the original study to normalize each 1D spatial expression data based on embryonic geometry derived from the image mask along the corresponding spatial axis and then log-transformed the normalized data (Junker et al. 2014). In the mouse olfactory mucosa data, we used the strategy described in (Segura et al. 2022) to initially fit 1D expression data for each gene along different spatial axes with polynomial fitting and then normalized them with the same procedure used for zebrafish embryo data.

For validation of the reconstructed zebrafish embryo data, we used an additional external Stereo-seq dataset that provides a spatiotemporal mapping of the zebrafish development that provides a more granular, ground-truth slice-wise view of gene expression Liu et al. (2022). We specifically used the embryo data at 5.25 hours post-fertilization (hpf) from this dataset to compare with our reconstruction on the tomo-seq data, which was in the shield stage (6 hpf). Though at slightly different time points, both are in the gastrula period of zebrafish embryo development Westerfield (2007).

### 4.8 Alignment of Stereo-seq and reconstructed RNA tomography slices

To use the slice-wise Stereo-seq zebrafish embryo data as a validation dataset to further assess reconstruction performance, twelve middle slices of the reconstructed gene expression data needed to be aligned with the twelve available Stereo-seq slices. Note, the reconstructed slices are 18*µm* and the Stereo-seq slices are 12*µm* in thickness. In order to do this, we first normalized the gene expression of the Stereo-seq data and the reconstructed gene expression data by dividing by the magnitude of the gene expression matrix. We then used PASTE’s `pairwise_align` function with default values for the parameters to align the slices after normalization Zeira et al. (2022). The output of the function is a list of the input slices–two in our case–with the coordinates of their spots adjusted so as to be more aligned with each other. We then use these aligned slices as input to our Moran score calculation.

### 4.9 Compared methods

CTFacTomo is benchmarked against the two available best methods, IPF algorithm (Fienberg 1970; Junker et al. 2014) and probabilistic graphical model Tomographer (Schede et al. 2021).

**IPF** is a simple scaling-based procedure for estimating the fitted tensor, 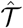, from the given marginals of the target tensor, 𝒯, which alternately minimizes the discrepancies between the marginals of 𝒯 and 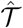 along different axes until convergence. In this study, we followed the MATLAB implementation described in Junker et al. (2014);

Segura et al. (2022) for 3D spatial transcriptome reconstruction with Tomo-seq (Junker et al. 2014) and rewrote the IPF algorithm in Python.

**Tomographer** is a compressed sensing algorithm to reconstruct the matrix from sampled marginal distributions by maximizing the posterior of their joint distribution. It is specifically designed for 2D spatial transcriptome reconstruction with STRP-seq (Schede et al. 2021), an RNA tomography technique that sections the tissue into consecutive slices and further cuts slices into strips at different angles. For 3D transcriptome reconstruction, Tomographer can be adapted by reconstructing a 2D spatial transcriptome for each tissue slice at a specific location along the z-axis at a time and subsequently stacking these reconstructions. In this study, we used the Tomographer package provided in the work (Schede et al. 2021) for our experiments.

Note that Tomographer was designed for 2D STRP-seq data, there is no 3D STRP-seq data for evaluation. Thus, the comparison to Tomographer is only conducted in the projected 3D ST datasets. To make the methods comparable, we applied Tomographer to the projected ST data with the secondary slices cut at 0 and 90 degrees.

## 5 Declarations

### Ethics approval and consent to participate

Not applicable.

### Consent for publication

The informed consent for all image data used in this study are obtained from the relevant organizations.

### Availability of data and materials

CTFacTomo is implemented in Python and the code is publicly available through GitHub at https://github.com/kuanglab/CTFacTomo. The datasets analyzed in this study are available in raw form from their original sources. Specifically, the 3D spatial transcriptomics data used to simulate RNA tomography are derived from either low-resolution ST1K or high-resolution Stereo-seq platforms, where data for the human heart tissue were obtained from the project website https://data.mendeley.com/datasets/mbvhhf8m62/2, mouse brain data were collected from https://molecularatlas.org/download-data, and drosophila embryo data are accessible at https://db.cngb.org/stomics/flysta3d/. Tomo-seq data for 3D spatial transcriptome reconstruction from the zebrafish embryo at the shield stage, along with the ISH images of marker genes, are available at https://www.ncbi.nlm.nih.gov/geo/query/acc.cgi?acc=GSE59873, while the Stereo-seq data of the zebrafish embryo at the shield stage are available at https://db.cngb.org/stomics/zesta/download/. The tomo-seq data from mouse olfactory mucosa and ISH images of marker genes can be accessed at https://www.ebi.ac.uk/biostudies/arrayexpress/studies/E-MTAB-10211.

### Authors’ contributions

R. Kuang designed the study; T. Song and R. Kuang implemented the algorithm, T. Song, Q. Nguyen, and C. Broadbent collected and analyzed data. T. Song, Q. Nguyen, and R. Kuang drafted the manuscript. All authors have read, edited, and approved the final manuscript.

## Acknowledgments

We sincerely thank Dr. Mayra Luisa, Dr. Antonio Scialdone, and Dr. Luis R. Saraiva for providing the mouse olfactory mucosa data and the scripts necessary to perform the experiments and the ISH images to evaluate the results.

