## Supplementary document for "CTFacTomo: Reconstructing 3D Spatial Structures of RNA Tomography Transcriptomes by Collapsed Tensor Factorization"

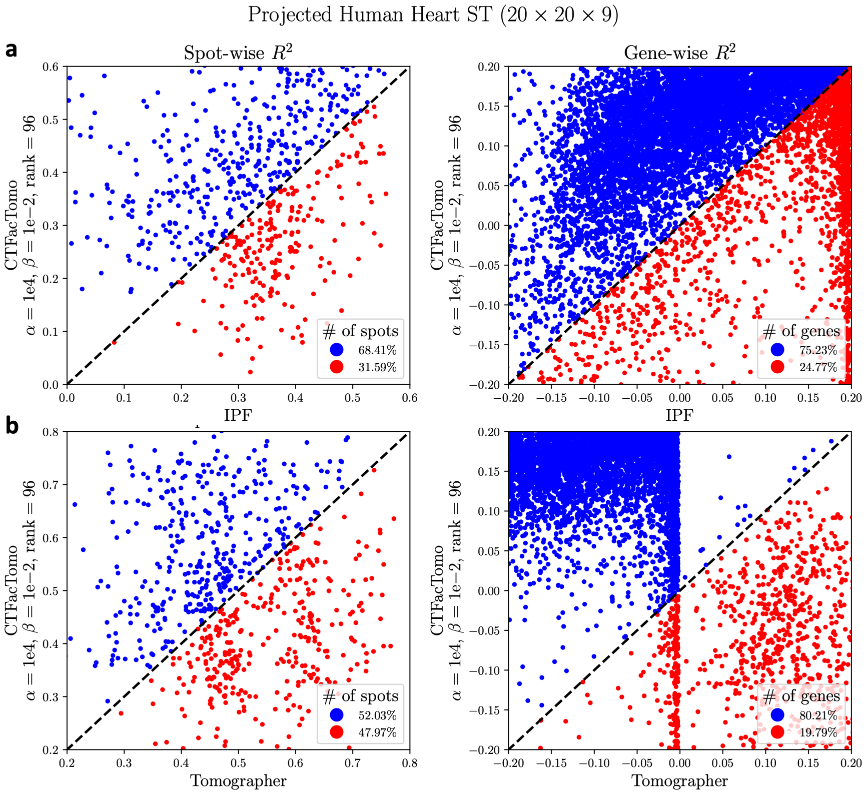


**Fig. S1. Comparison of Reconstruction Performance on the Projected Human Heart data**. **a** IPF (x-axis) and CTFacTomo (y-axis) **b** Tomographer (x-axis) and CTFacTomo (y-axis) ***left*** Scatter plot of spot-wise $R^2$ from different methods; ***right*** Scatter plot of gene-wise $R^2$ from different methods. Each dot represents either a spot or a gene, and blue dots in both scatter plots indicate either the reconstructed expressions of spots or genes from CTFacTomo achieving higher $R^2$ compared to either IPF or Tomographer while red dots denote the reconstructed expressions of spots or genes from either IPF or Tomographer outperform CTFacTomo in terms of $R^2$.

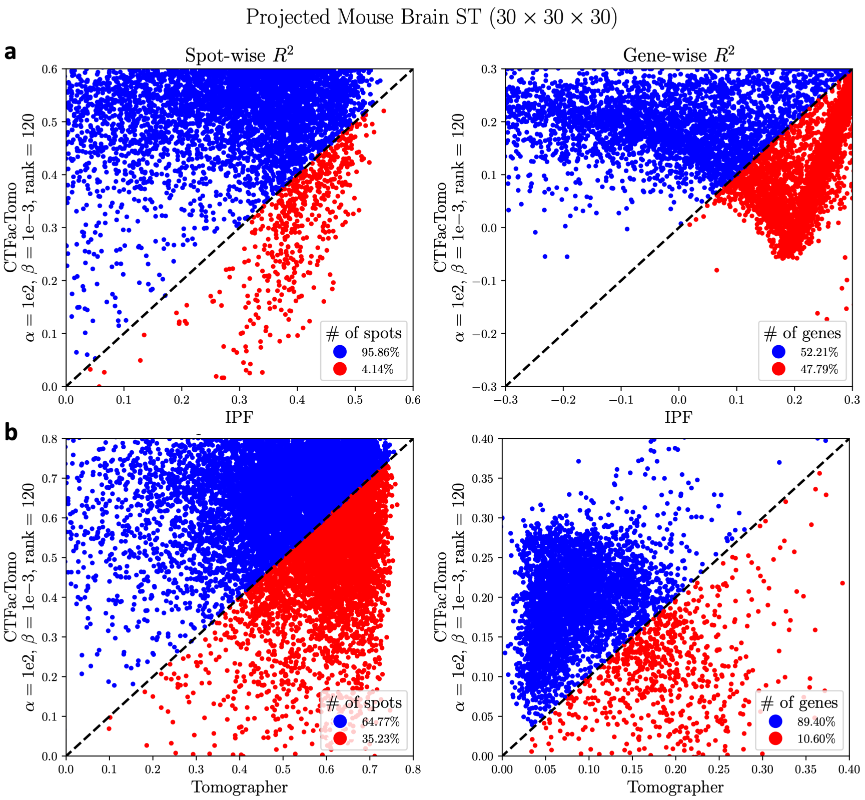


**Fig. S2. Comparison of Reconstruction Performance on the Projected Mouse Brain data**. **a** IPF (x-axis) and CTFacTomo (y-axis) **b** Tomographer (x-axis) and CTFacTomo (y-axis) ***left*** Scatter plot of spot-wise $\mathrm{R}^2$ from different methods; ***right*** Scatter plot of gene-wise $\mathrm{R}^2$ from different methods. Each dot represents either a spot or a gene, and blue dots in both scatter plots indicate either the reconstructed expressions of spots or genes from CTFacTomo achieving higher $\mathrm{R}^2$ compared to either IPF or Tomographer while red dots denote the reconstructed expressions of spots or genes from either IPF or Tomographer outperform CTFacTomo in terms of $\mathrm{R}^2$.

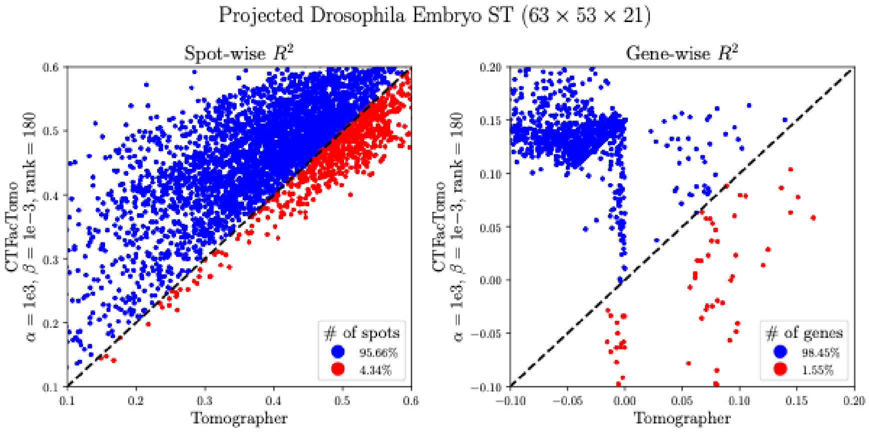


**Fig. S3. Comparison of Reconstruction Performance on the Projected Drosophila Embryo data**. ***left*** Scatter plot of spot-wise $R^2$ from Tomographer (x-axis) and CTFacTomo (y-axis); ***right*** Scatter plot of gene-wise $R^2$ from Tomographer (x-axis) and CTFacTomo (y-axis). Each dot represents either a spot or a gene, and blue dots in both scatter plots indicate either the reconstructed expressions of spots or genes from CTFacTomo achieving higher $R^2$ compared to Tomographer while red dots denote the reconstructed expressions of spots or genes from IPF outperform either Tomographer in terms of $R^2$.

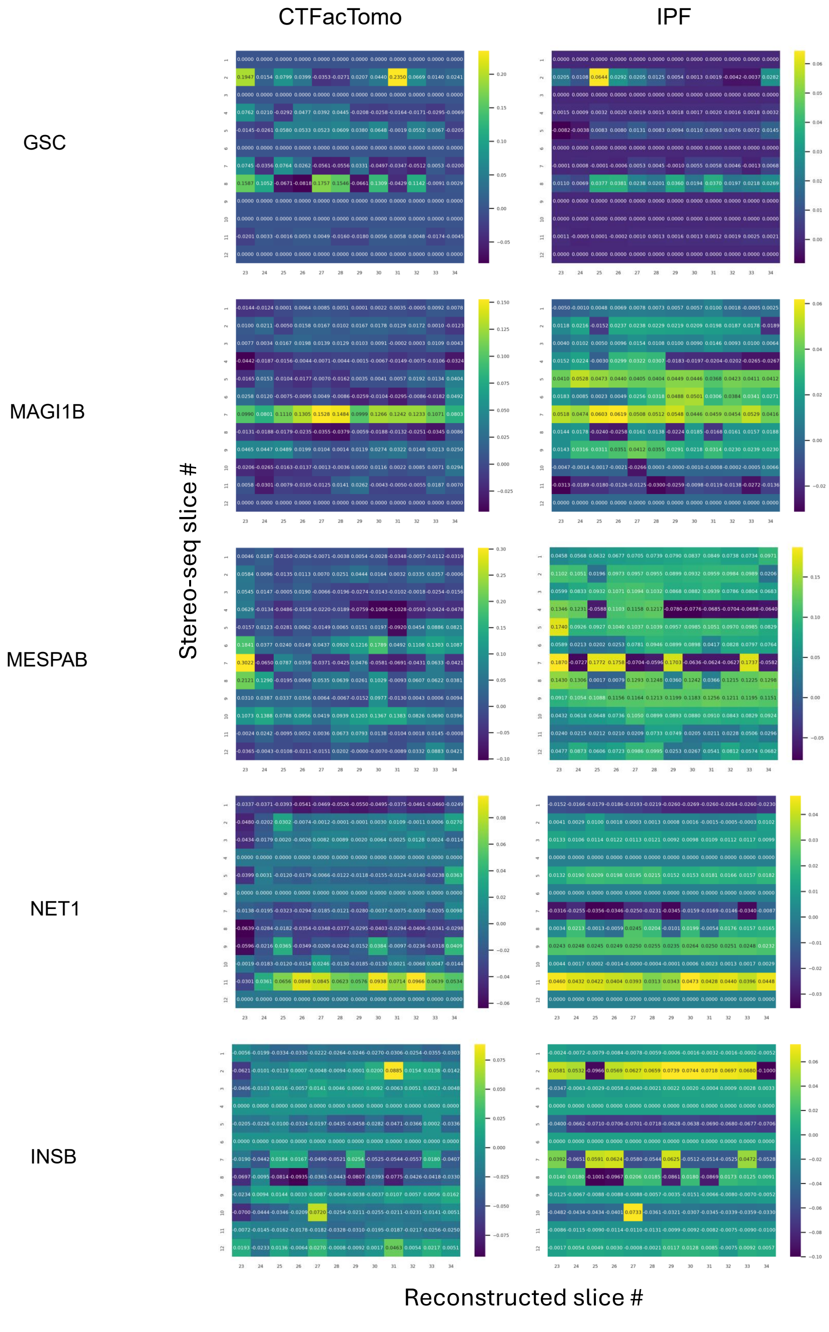


**Fig. S4. Moran scores between aligned slices for Stereo-seq validation.** Heatmap visualizations for the Moran scores between the reconstructed slices and Stereo-seq slices for five marker genes (*GSC*, *MAGI1B*, *MESPAB*, *NET1*, *INSB*). ***left*** Scores when comparing CTFacTomo reconstructed slices with Stereo-seq slices. ***right*** Scores when comparing IPF reconstructed slices with Stereo-seq slices. All spots within the reconstructed slice were considered. CTFacTomo always has a higher max score when comparing the heatmaps.

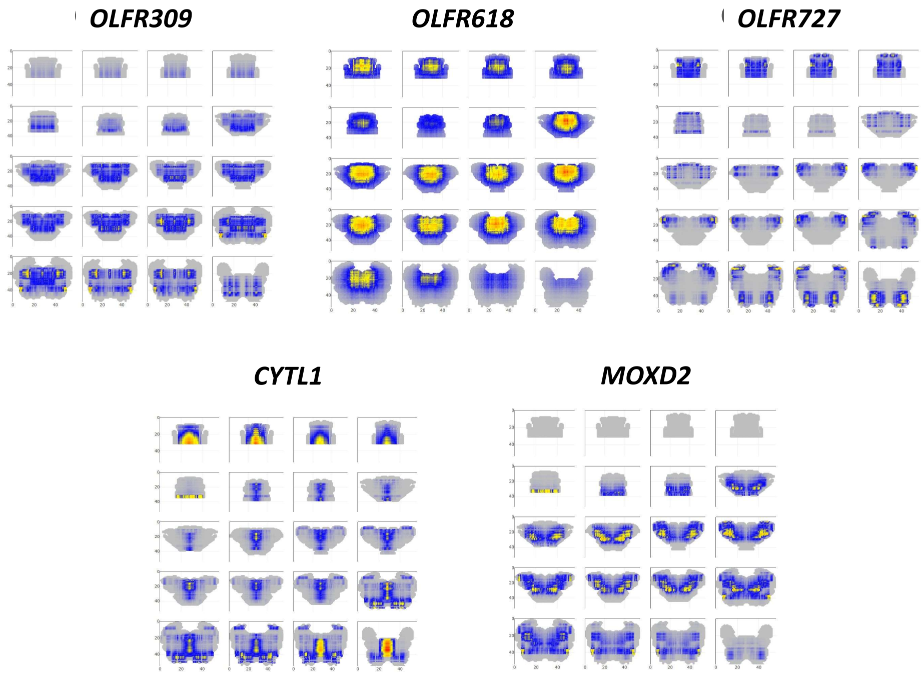


**Fig. S5. Visualizations for the CTFacTomo Reconstruction of the Mouse Olfactory Mucosa on Five Marker Genes.** The visualized slices are along the anteroposterior axis. The color scale ranges from blue to red, indicating low to high levels of relative expression for that gene, respectively. Gray indicates no expression.

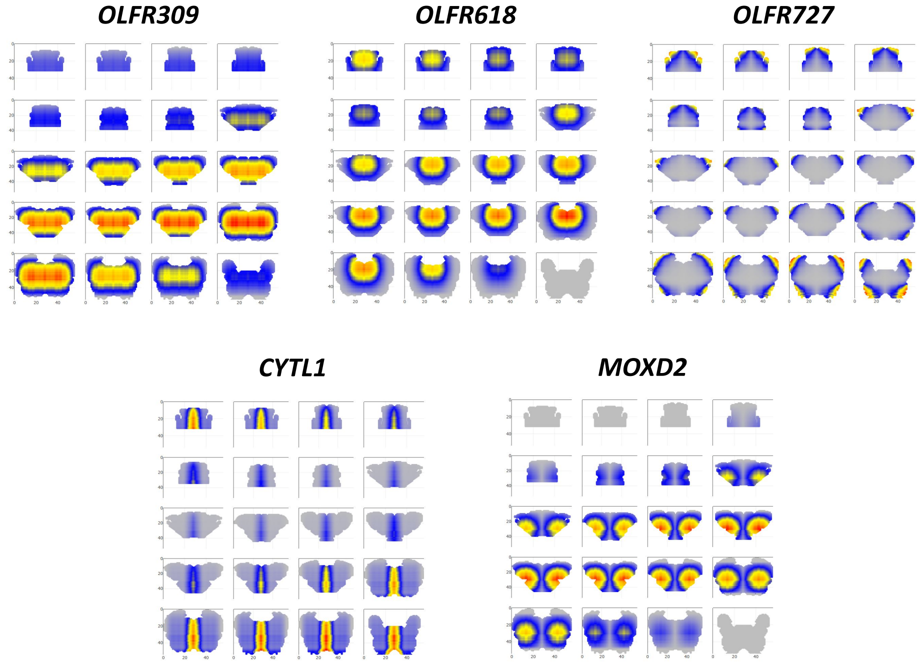


**Fig. S6. Visualizations for the IPF Reconstruction of the Mouse Olfactory Mucosa on Five Marker Genes.** The visualized slices are along the anteroposterior axis. The color scale ranges from blue to red, indicating low to high levels of relative expression for that gene, respectively. Gray indicates no expression.
